## Supplementary material for "Tracing the function expansion for a primordial protein fold in the era of fold-based function prediction: β-trefoil"

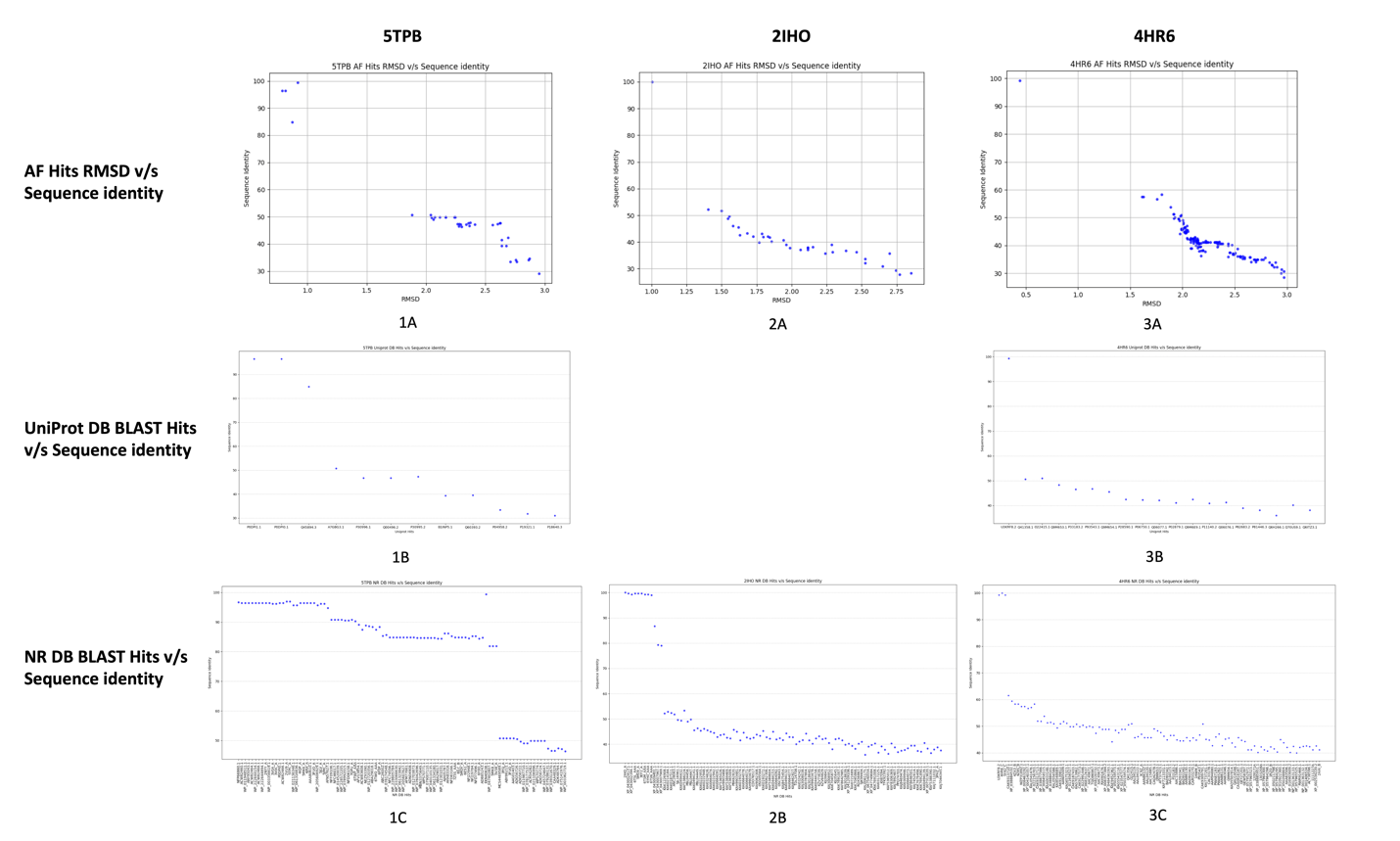

**S1 Fig: For the PDB structures from table 1 with gap between low r.m.s.d. (<1.5) and high sequence identity (> 50% ): 1A, 2A and 3A show r.m.s.d. (x-axis) v/s Sequence identity (y-axis) plots for the AFdb Hits from 5TPB, 2IHO and 4HR6 respectively (first row), 1B and 3B show UniProt database BLAST Hits (x-axis) v/s Sequence identity (y-axis) for 5TPB and 4HR6 respectively (second row) and 1C, 2B and 3C show NR database BLAST Hits (x-axis) v/s Sequence identity (y-axis) for 5TPB, 2IHO and 4HR6 respectively (third row)**

**A**

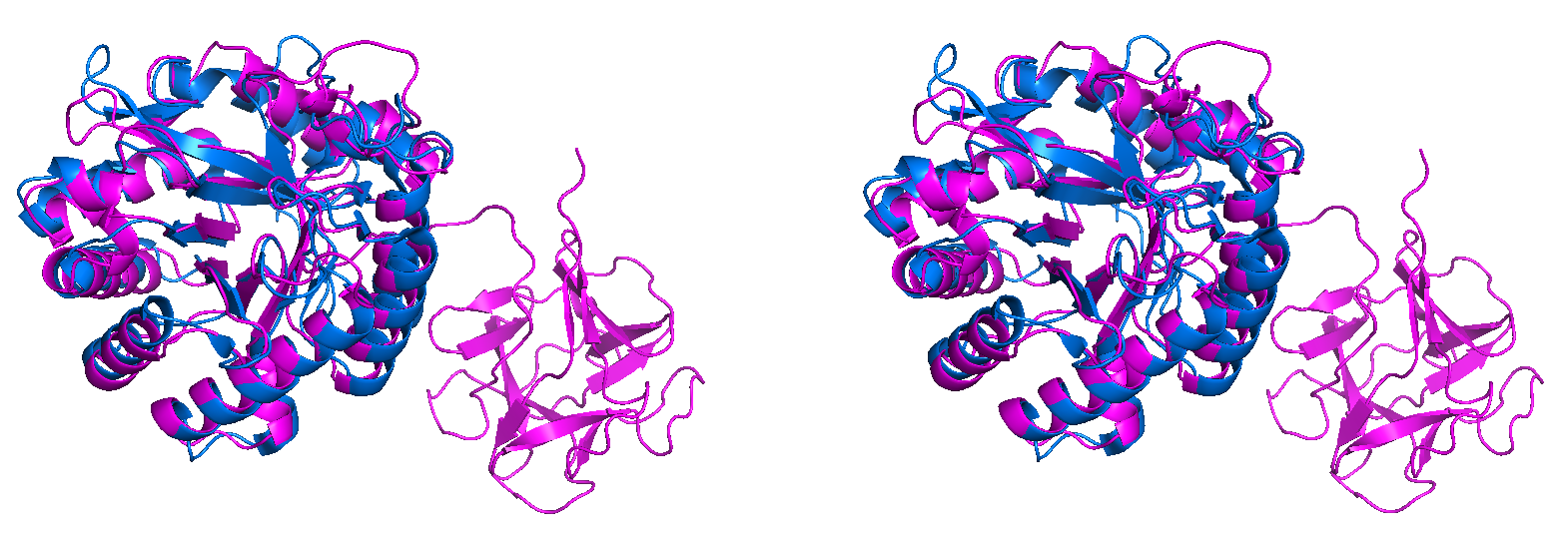

A0A6I5GSK5 (PMID: [29146152](https://pubmed.ncbi.nlm.nih.gov/29146152/)) sharing its chitinase domain with 2DSK from *Pyrococcus furiosus* with an RMSD of 1.342 Å (**PMID:** [**17183162**](https://www.rcsb.org/search?q=rcsb_pubmed_container_identifiers.pubmed_id:17183162)) (Figure 2a)

**B**

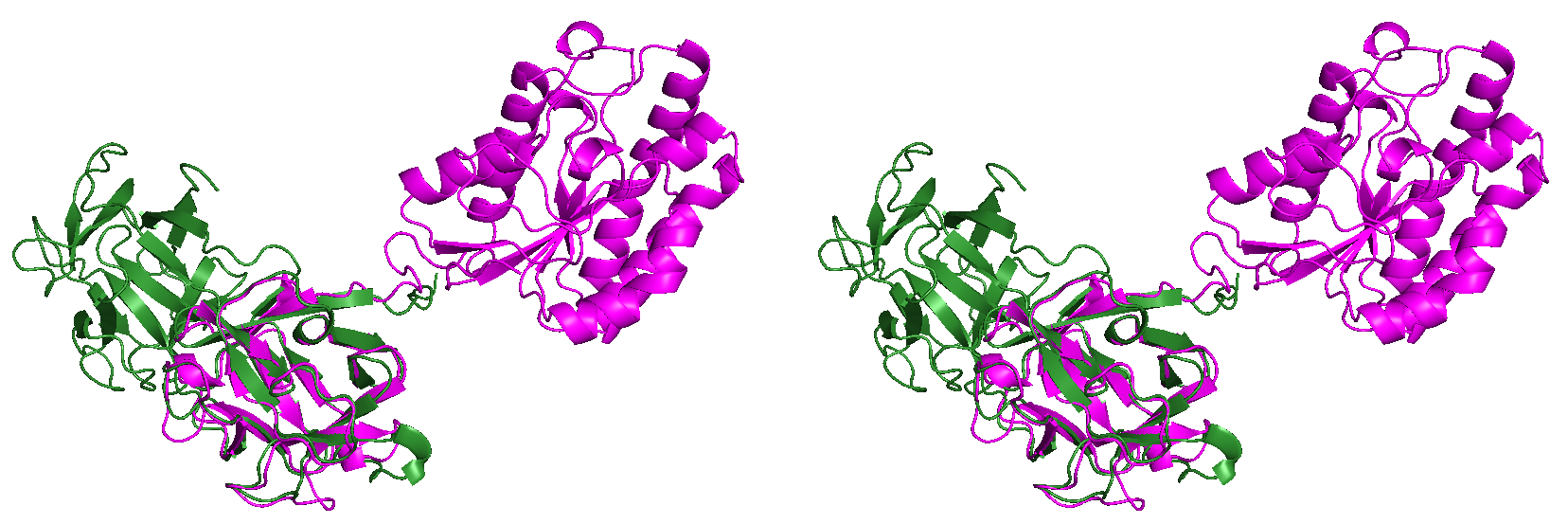

A0A318NTU0 giving an RMSD of 0.795 Å against the β-trefoil domain of 1HWM (Figure 2b)

**C**

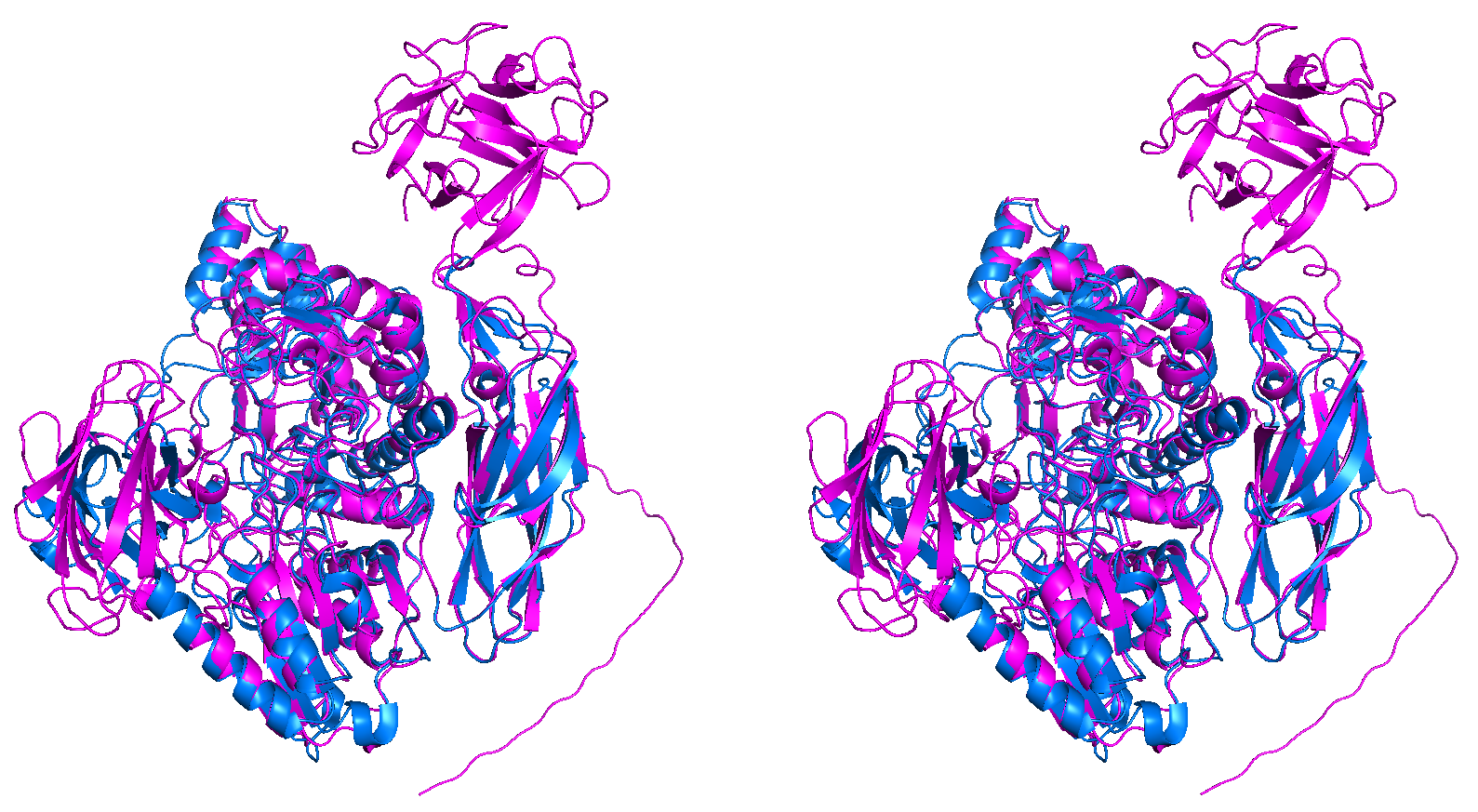
A0A4Q2KX23 (**PMID**: not available) sharing its beta-glucosidase domain with 4I3G from *Streptomyces venezuelae* with an RMSD 1.016 Å (**PMID** [**23225731**](https://www.rcsb.org/search?q=rcsb_pubmed_container_identifiers.pubmed_id:23225731))

**D**

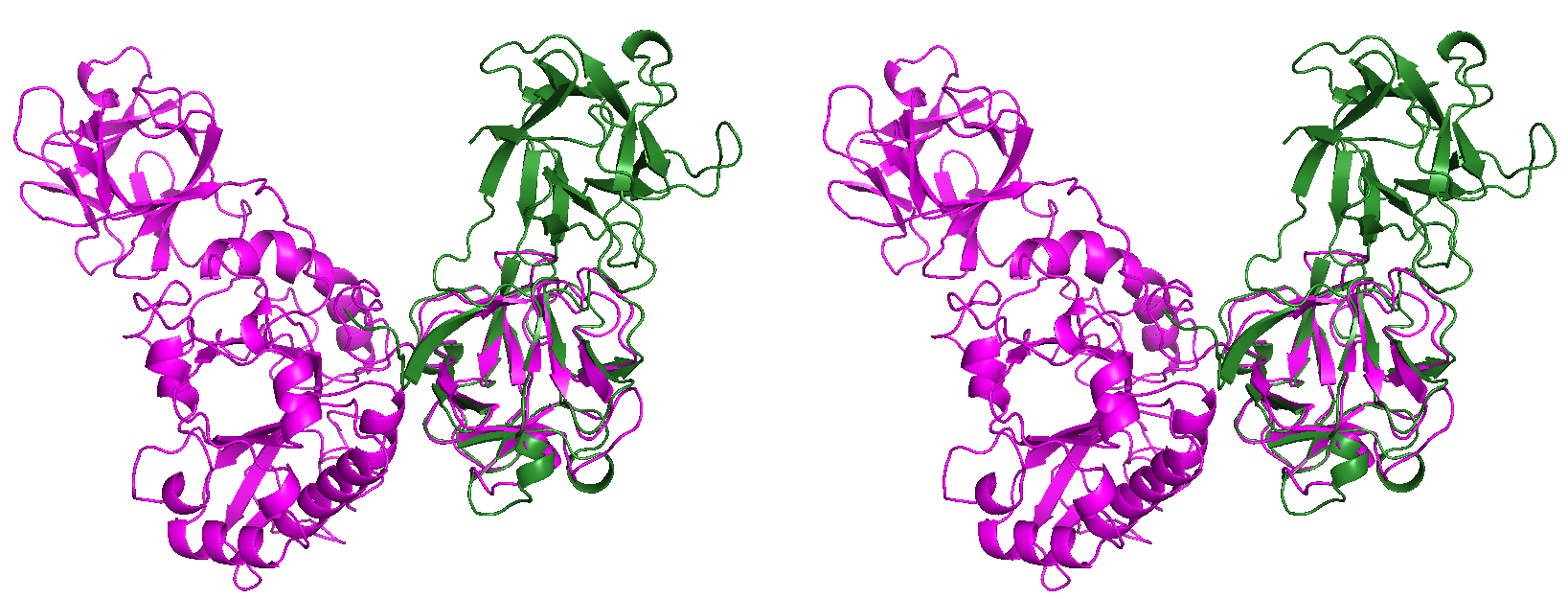

A0A1I6E245 (**PMID**: Not published yet) giving an RMSD of 1.867 Å against the β-trefoil domain of 4HR6

**E**

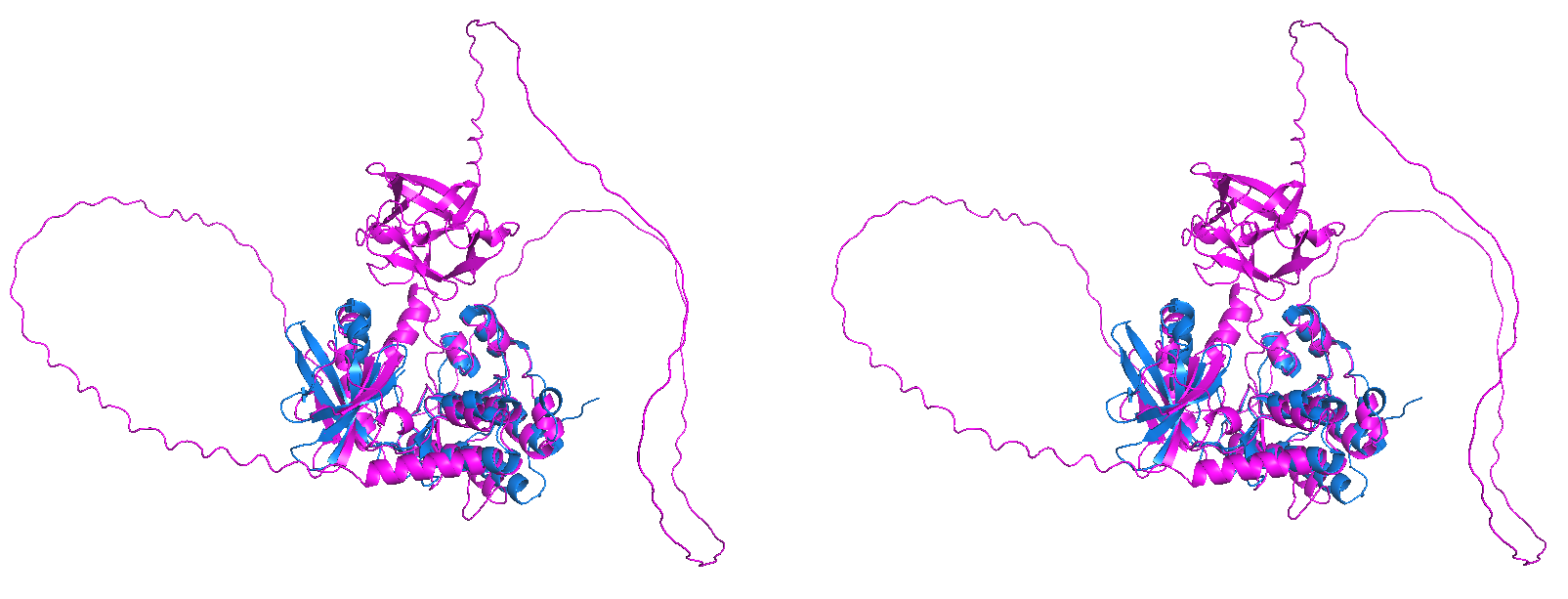

A0A3D9WMQ9 (**PMID:** not available) sharing its protein kinase with 3F61 from *Mycobacterium tuberculosis* with an RMSD of 1.95 Å

**F**

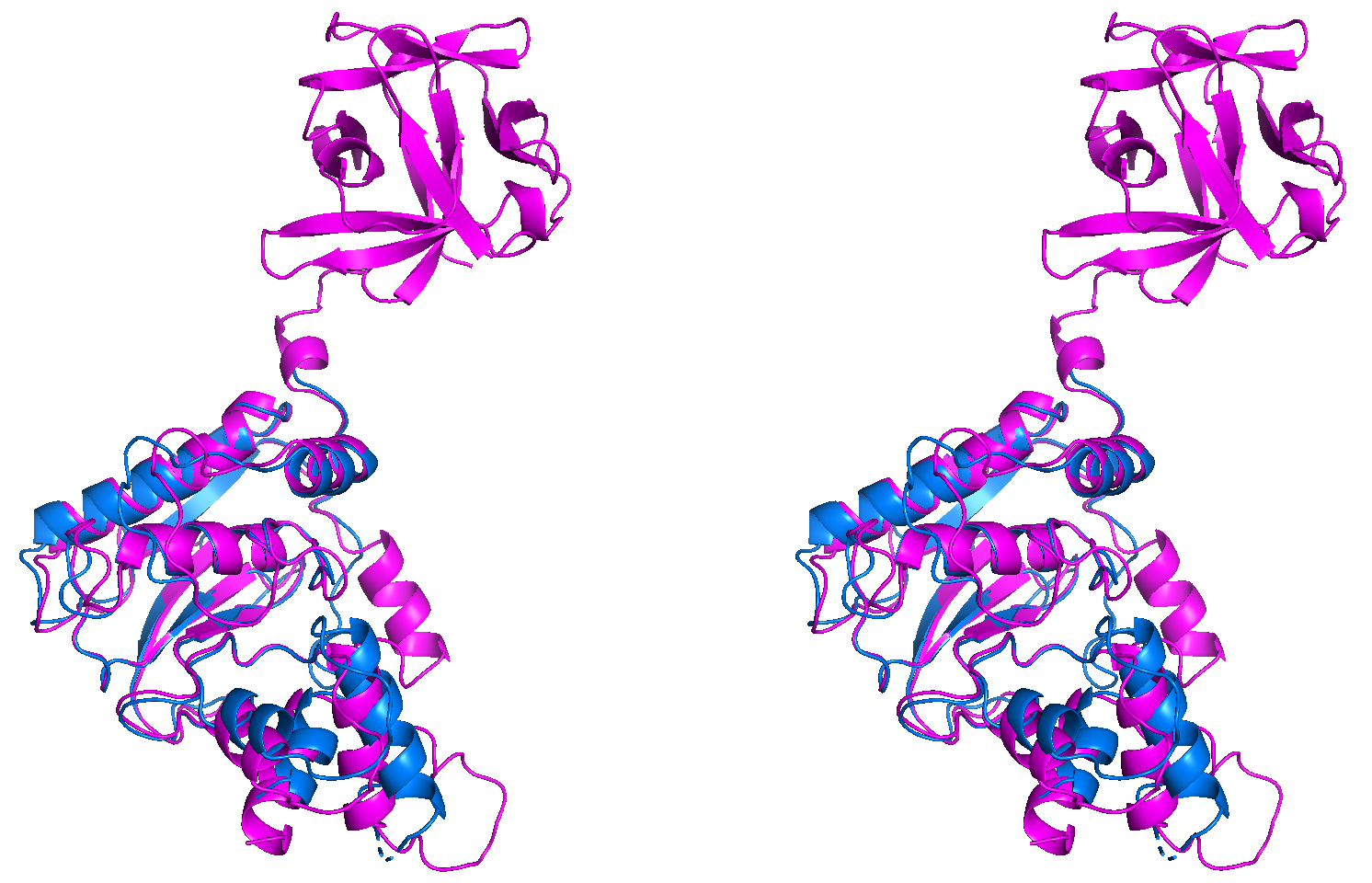

A0A820M8C8 (**PMID**: not available) sharing its metallopeptidase domain with 1SLM from *Homo sapiens* with an RMSD of 0.824 Å

**S2 Fig: Characterized hits with diverse and novel domain architecture superimposed with PDB - enlarged view of each superimposed pair as shown in Figure 3, in the same order**

**A**

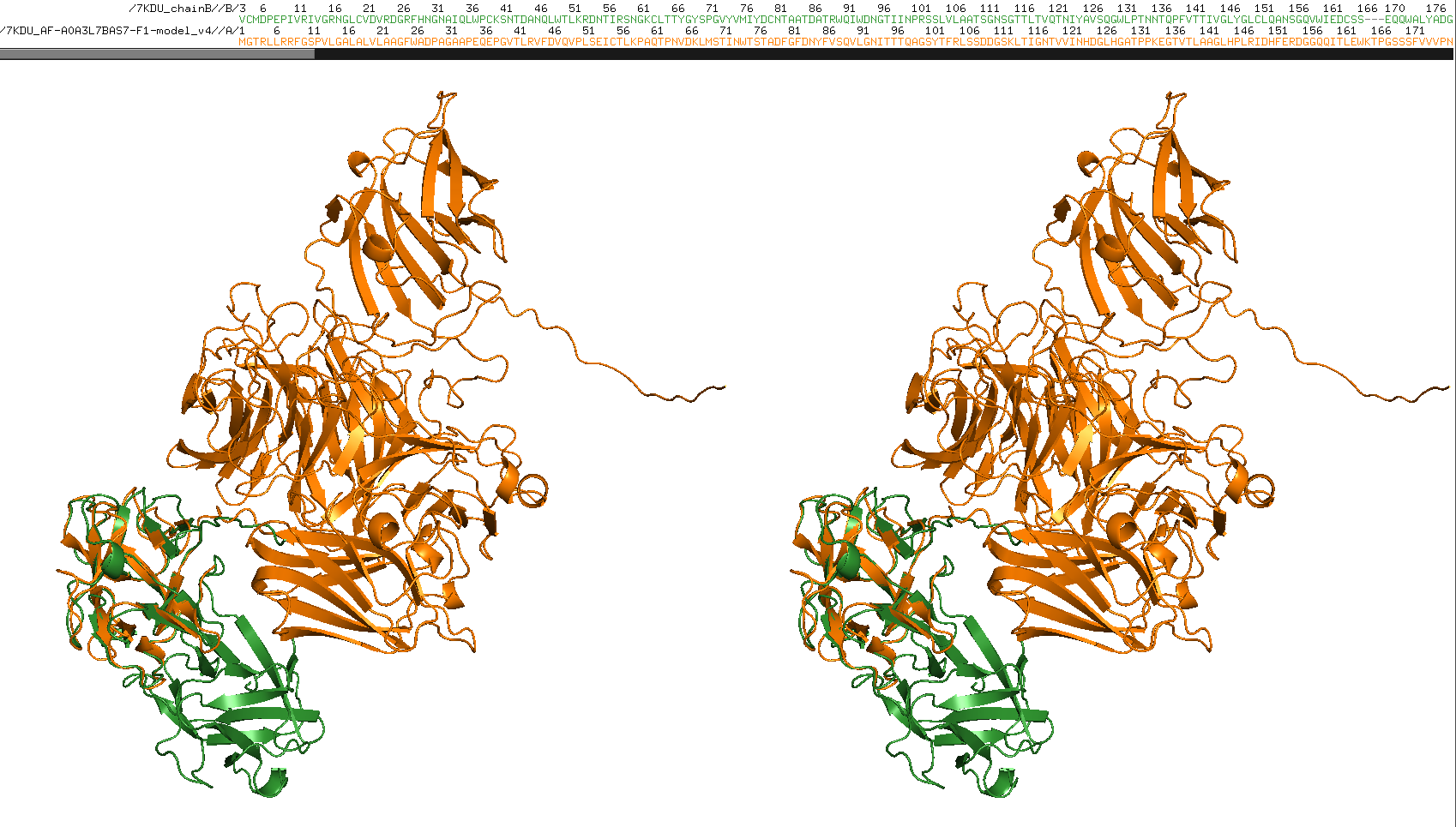

7KDU (Ricin) against A0A3L7BAS7

RMSD: 0.686 Å

**B**

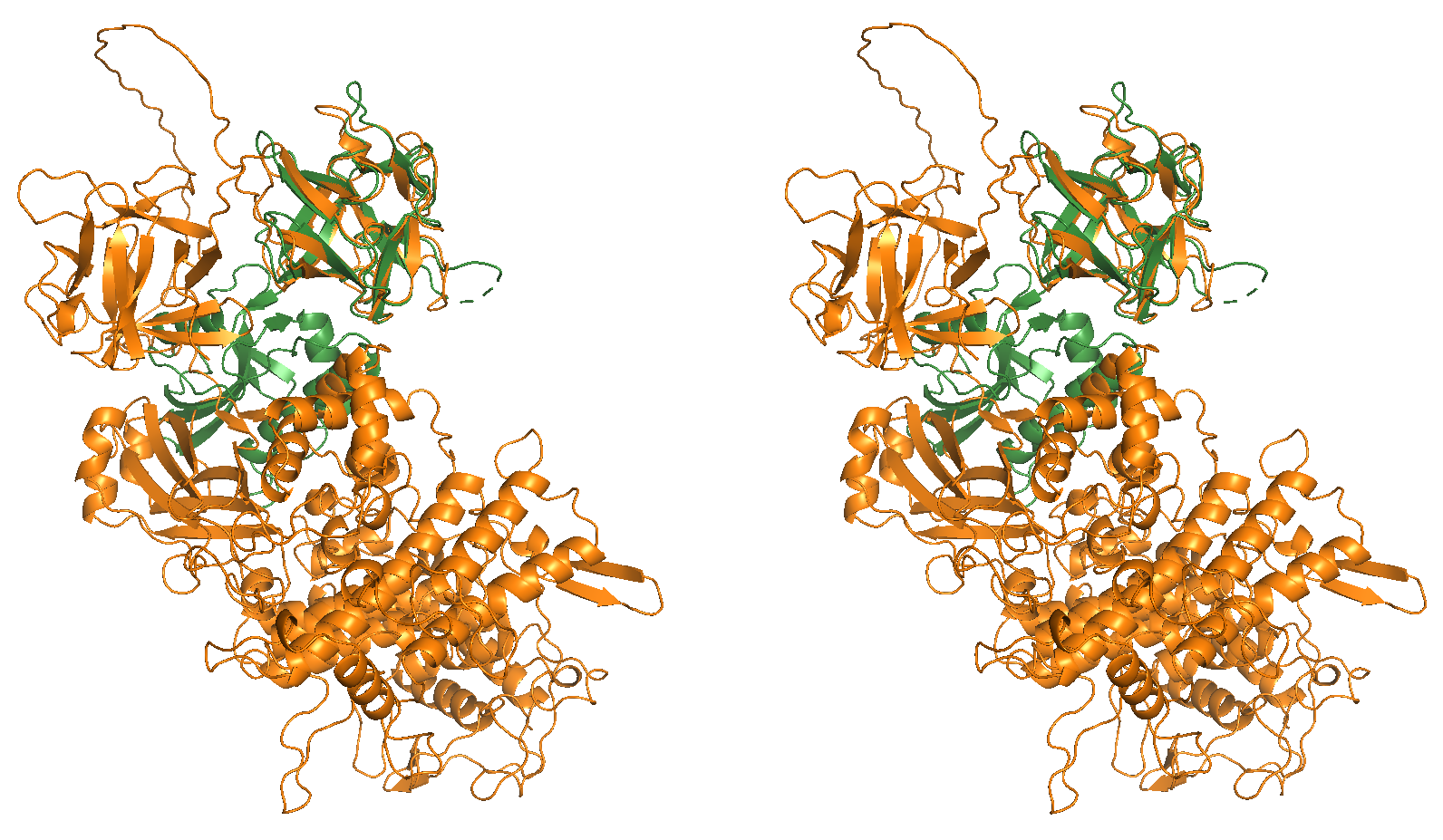

3PHZ (Ricin) against A0A0U3MKC8

RMSD: 0.888 Å

**C**

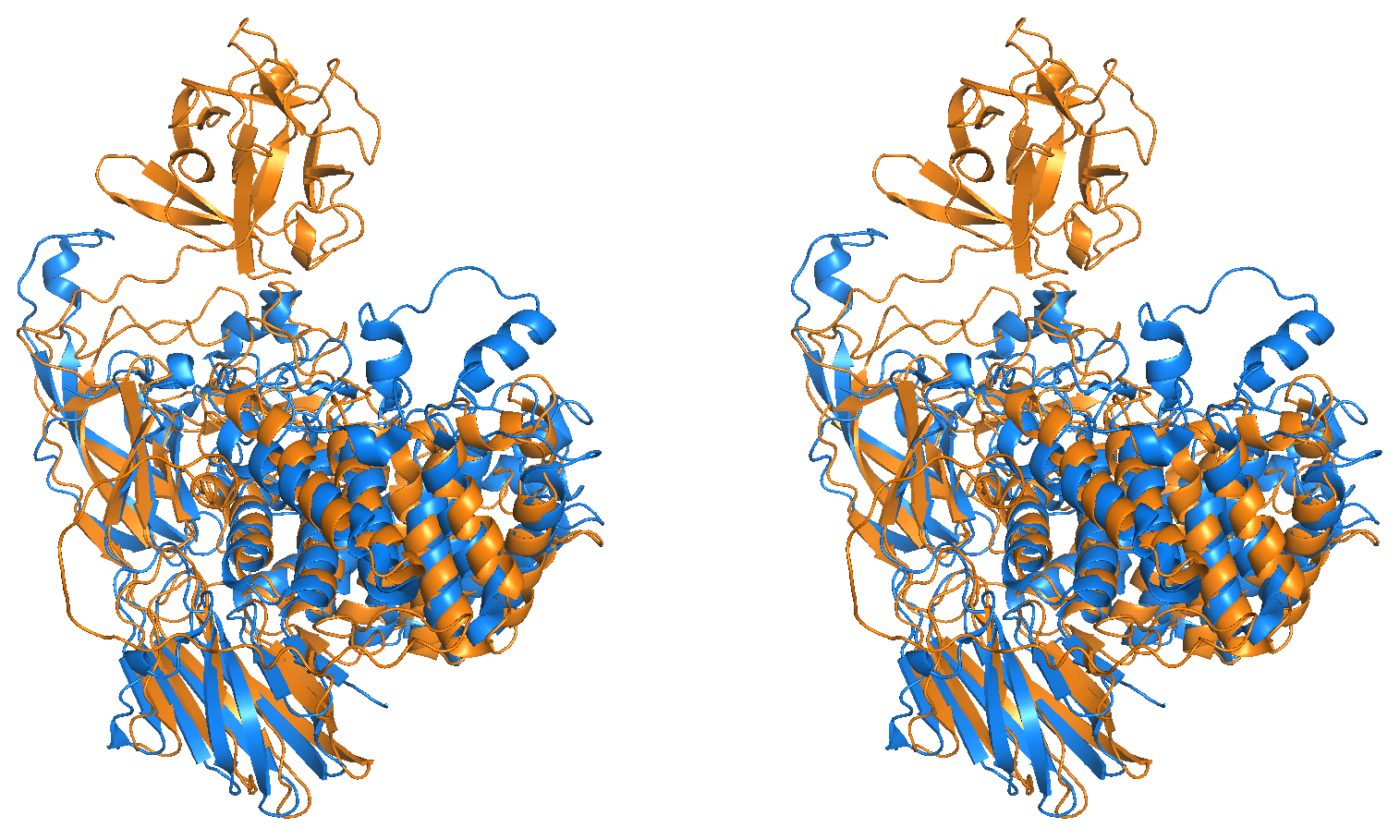

A0A191US67 sharing the Beta-L- arabinofuranosidase domain with 4QJY (Hydrolase- GH127 Beta-L-arabinofuranosidase from *Geobacillus stearothermophilus*)

RMSD: 2.83 Å

**D**

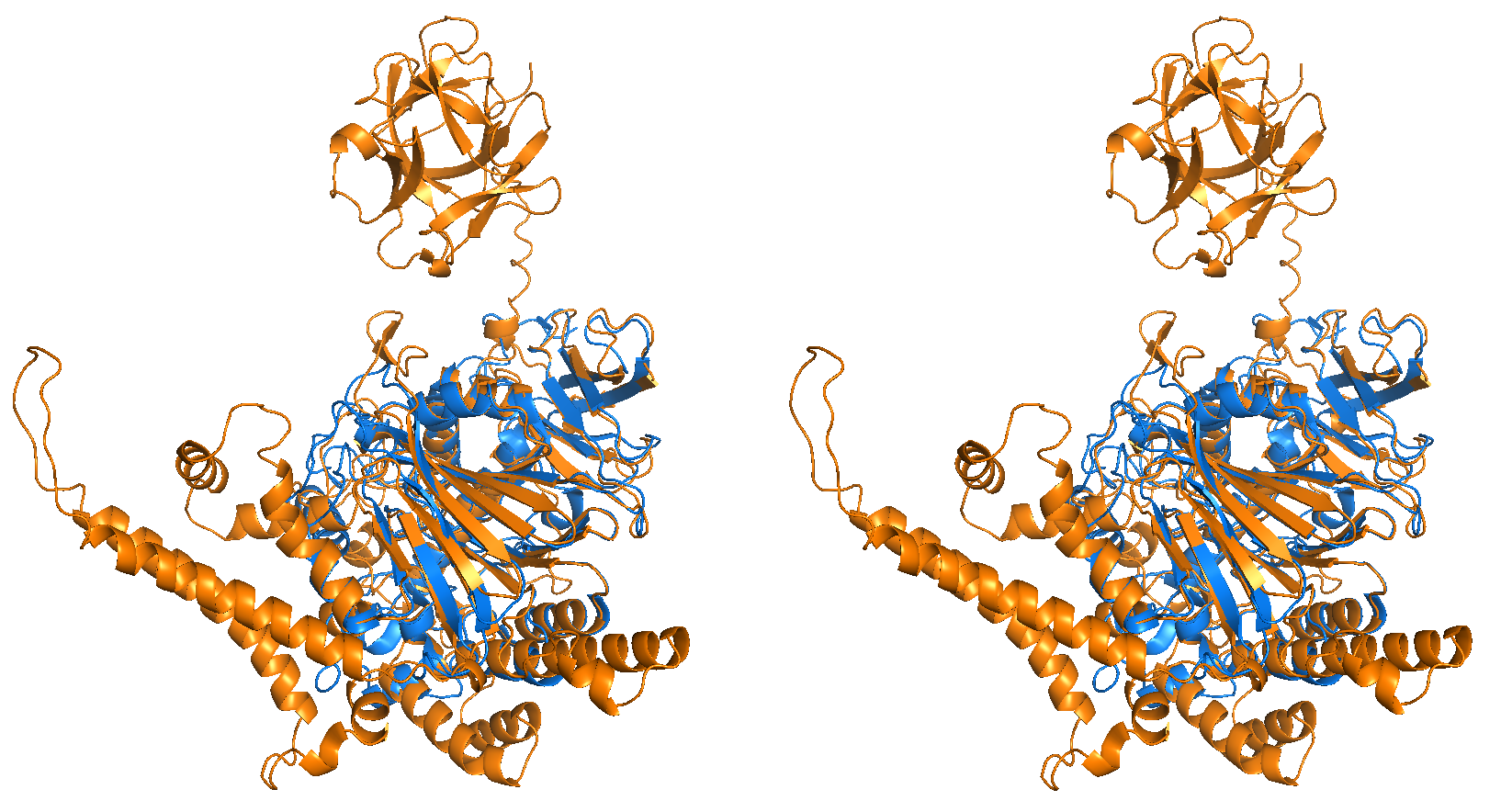

A0A844M7R0 sharing the PhoD domain with 2YEQ (Hydrolase- PhoD from *Bacillus subtilis*)

RMSD: 0.741 Å

**E**

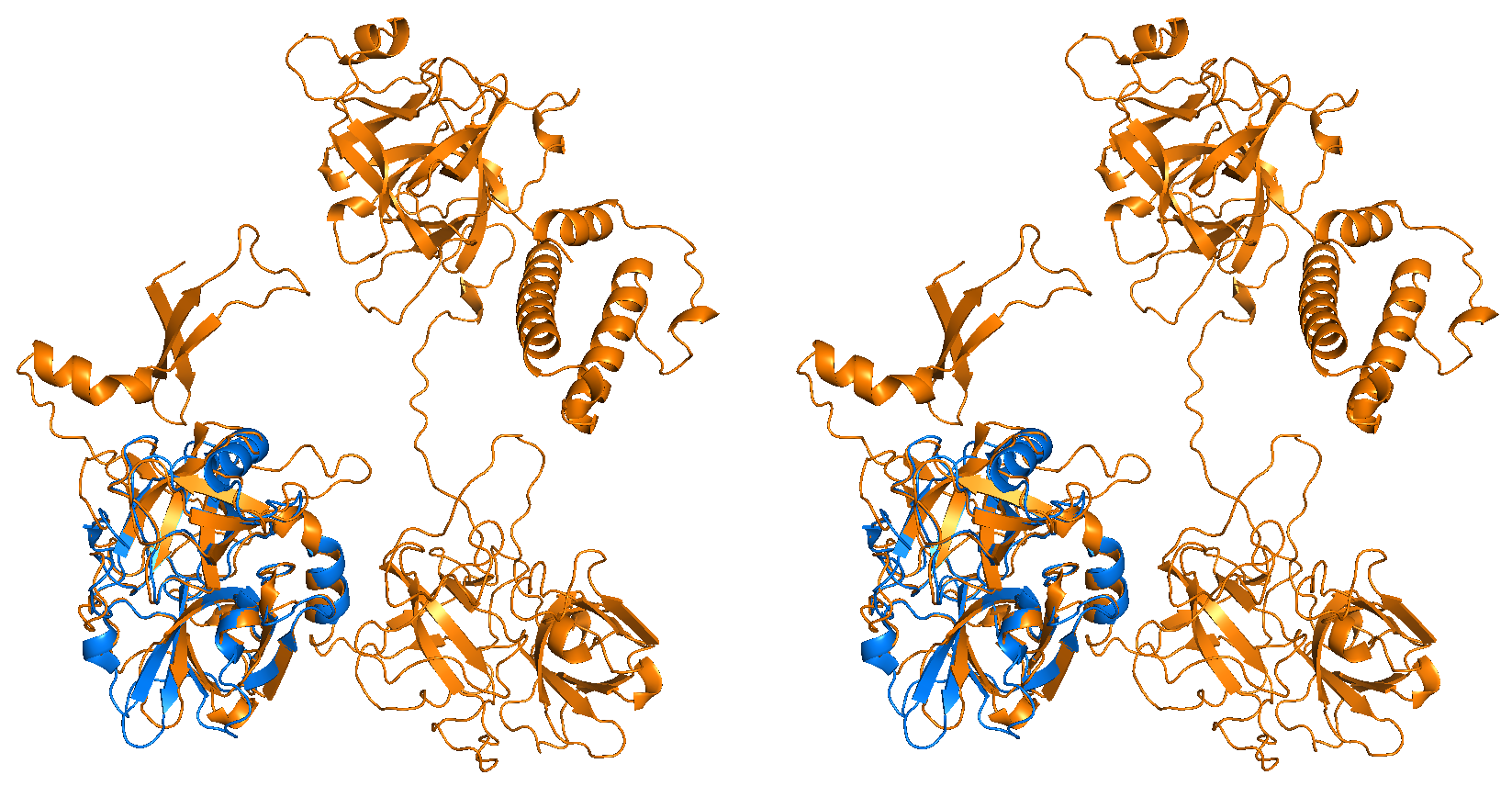

A0A820T4Y8 sharing the Trypsin domain with 7N7X (plasma kallikrein domain that contains trypsin domain from *Homo sapiens*)

RMSD: 0.628

**F**

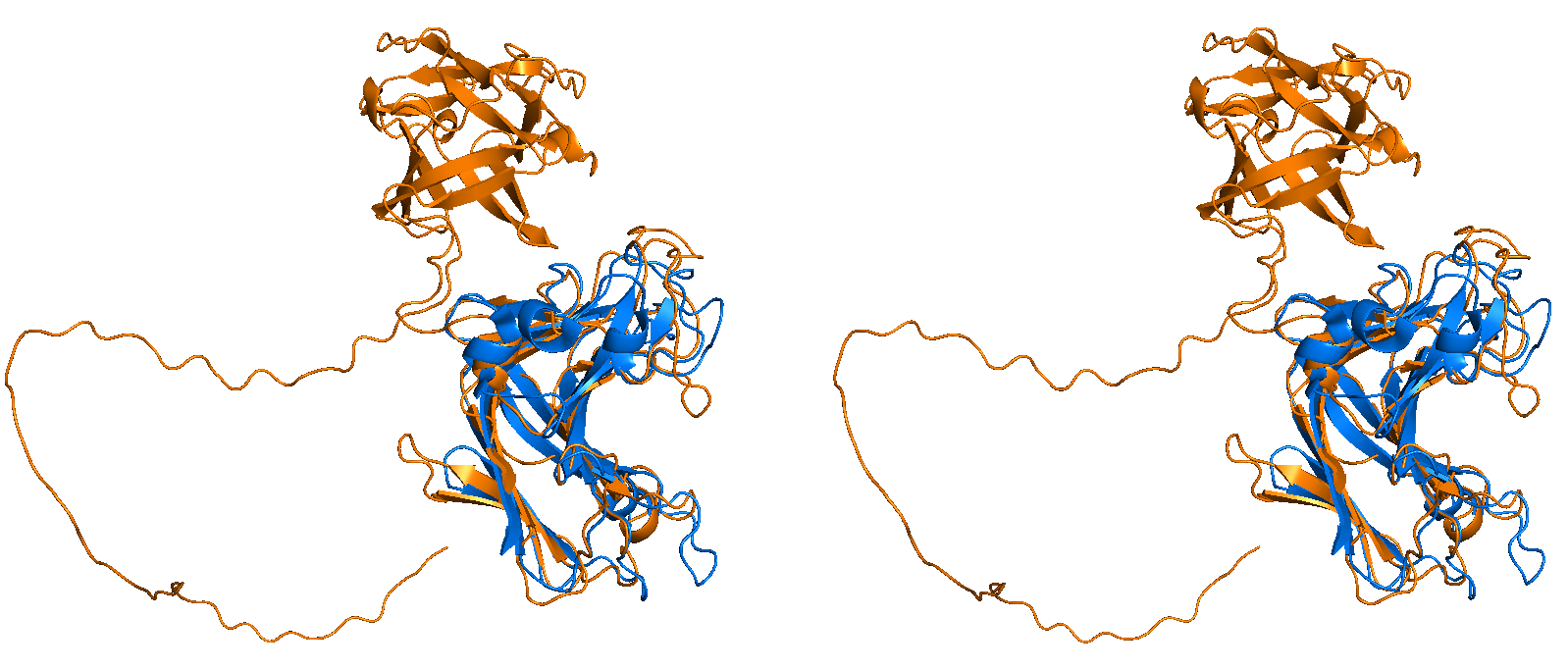

A0A3A9W2R1 sharing the Alginate lyase domain with 3ZPY (Alginate lyase from *Zobellia galactanivorans* **PMID**: [**36548760**](https://www.rcsb.org/search?q=rcsb_pubmed_container_identifiers.pubmed_id:36548760))

RMSD: 0.976 Å

**S3 Fig: Uncharacterized hits with diverse and novel domain architecture superimposed with PDB - enlarged view of each superimposed pair as shown in Figure 5, in the same order**

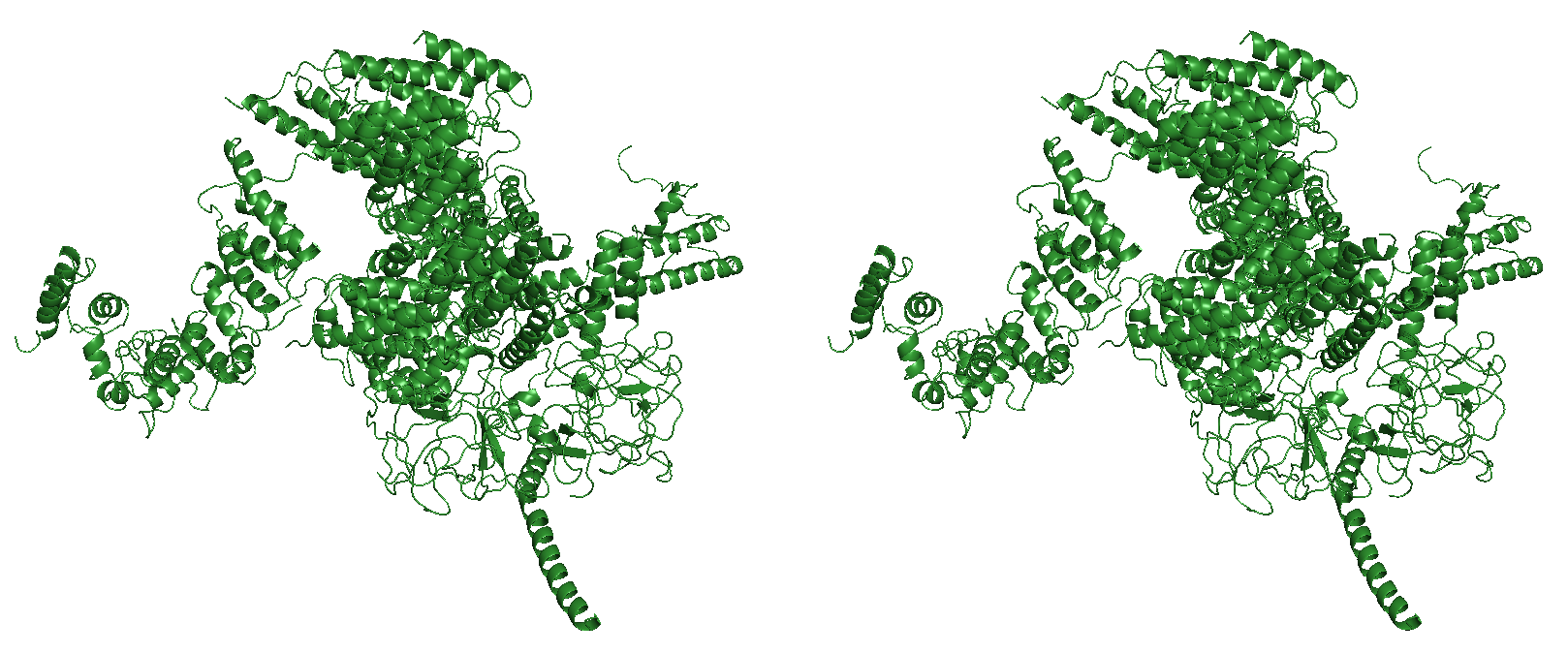

**S4 Fig:**

6MU1 (Inositol 3-P receptor)

[Note: β-trefoil domains (in this case, mostly without well-defined secondary structure elements) cover only 15% of the total structure which is as long as 2732 residues, therefore, the 30% and above coverage rule for filtering AFdb hits does not apply to this]

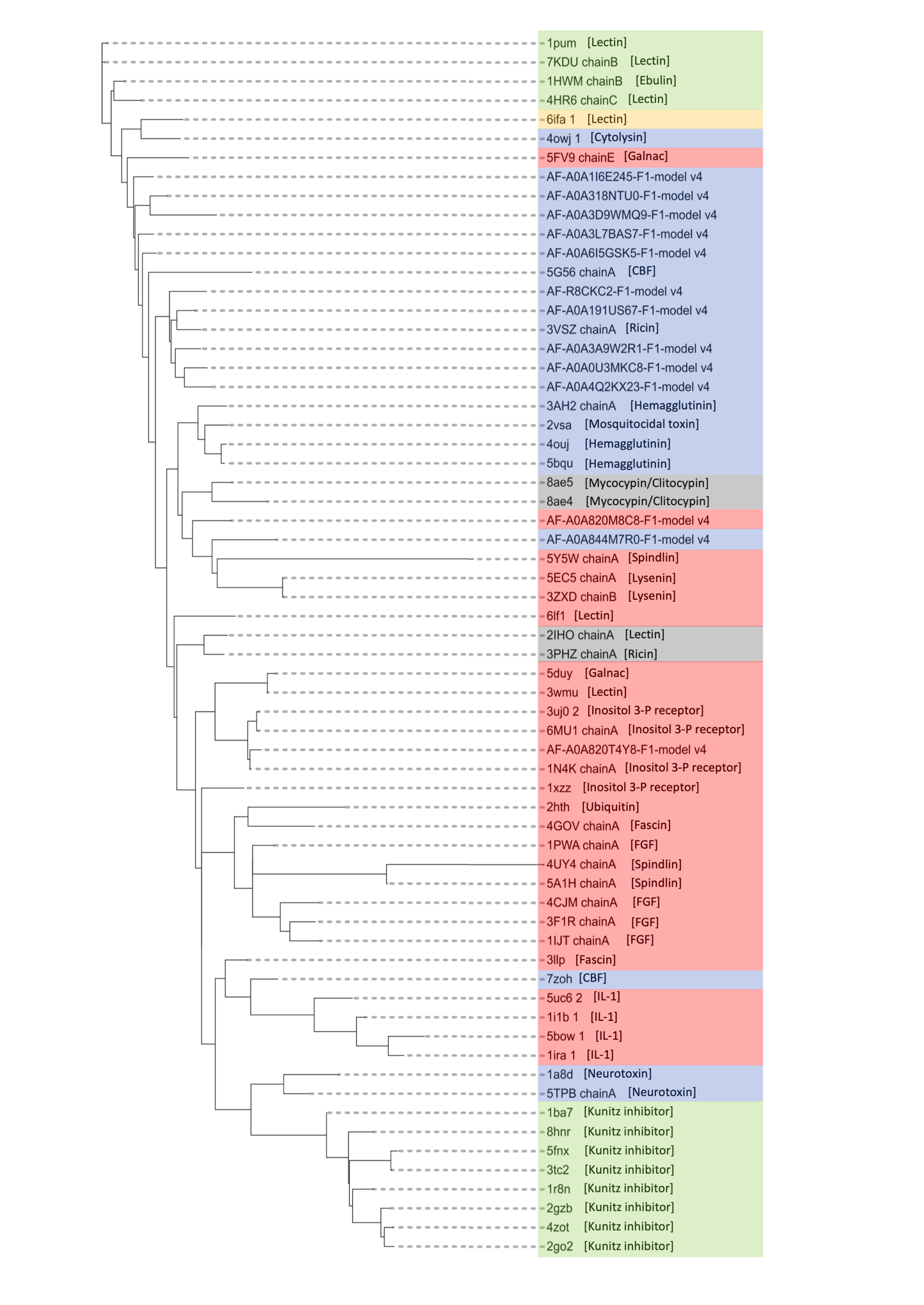

**S5 Fig: Phylogenetic tree obtained by structure-based sequence alignment of selected β-trefoil domains (filtered to include only those positions with less than 10% gaps) for all 64 structures from PDB and AFdb**

**S1 Table:** Novel domain architecture for proteins from AFdb containing trefoil fold

| **Uniprot ID** | **Organism** | **Novel Domain name** | **Domain Architecture** | **Remarks** |
| --- | --- | --- | --- | --- |
| A0A6I5GSK5  (Chitinase) | *Streptomyces sp. SID89*  *(Culturable bacteria)* | Glycosyl hydrolases family 18 | 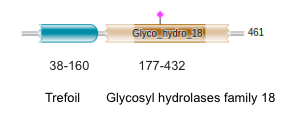 | Category-2  Glyco hydrolase family 43 found in Sundial (3VSZ)- Different fold compared to this |
| A0A318NTU0  (Lipase) | *Micromonospora arborensis* | Lipase | 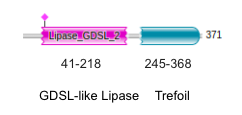 | Category-1 |
| A0A4Q2KX23 (Beta-glucosidase) | *Agromyces albus* | Glycosyl hydrolases family 3, PA14 and Fibronectin type- III | 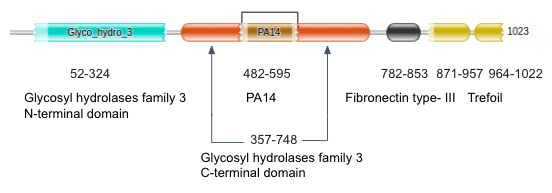 | Category-2 |
| A0A1I6E245  (Ricin-type β-trefoil lectin domain-containing protein) | *Lentzea waywayandensis* | Unknown domain | 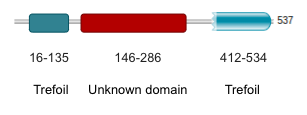 | Category-1 |
| A0A3D9WMQ9  (Serine/threonine protein kinase) | *Mycobacterium tuberculosis* | Protein kinase | 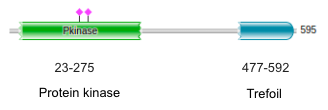 | Category-2 |
| A0A820M8C8 (Peptidase metallopeptidase domain-containing protein) | *Rotaria sp.*  *(Metazoa)* | Peptidoglycan-binding domain, Peptidase matrixin | 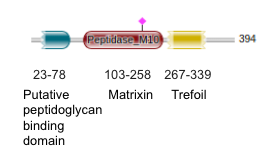 | Category-2 |
| A0A3L7BAS7 | *Micromonospora sp. CV4 (Culturable Bacteria)* | Alpha galactosidase A [N- terminal] | 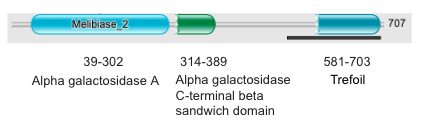 | Category-1 |
| A0A191US67 | *Streptomyces parvulus (Culturable bacteria)* | Beta-L- arabinofuranosidase, GH127 [N- terminal] | 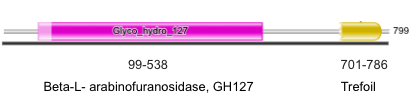 | Category-2 |
| A0A844M7R0 | *Scytonema sp. UIC 10036 (Culturable cyanobacteria)* | PhoD- like phosphatase (2 domains) [C- terminal] | 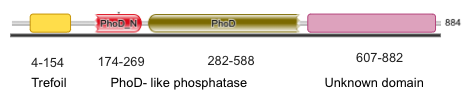 | Category-2 |
| R8CKC2 | *Bacillus cereus*  *(Culturable bacteria)* | Insecticidal Crystal Toxin, P42 [C- terminal] | 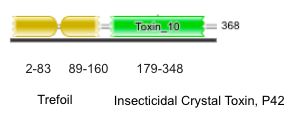 | Category-3 |
| A0A820T4Y8 | *Rotaria sp. Silwood2 (Metazoa)* | Trypsin (2 domains) [N- terminal] | 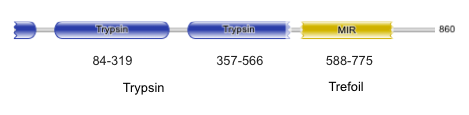 | Category-2 |
| A0A0U3MKC8 | *Roseateles depolymerans*  *(Culturable bacteria)* | Unknown domains [C- terminal] | 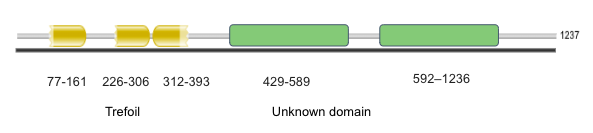 | Category-1 |
| A0A3A9W2R1 | *Aquimarina sp. BL5* | Alginate lyase | 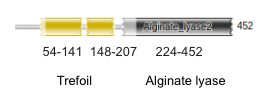 | Category-2 |
